# When “Noise” Isn’t Simply Noise: Deterministic Postural Drive During Noisy Galvanic Vestibular Stimulation (nGVS)

**DOI:** 10.64898/2026.04.20.719310

**Authors:** Dominique Rice, Kelci B. Hannan, Malynn Ewer, Christopher J Dakin

## Abstract

Age- and disease-related vestibular decline can cause dizziness and postural instability, motivating interventions such as noisy galvanic vestibular stimulation (nGVS). nGVS is commonly delivered at “subsensory” amplitudes and explained by stochastic resonance, yet because galvanic stimulation directly modulates vestibular afferents, even imperceptible currents may also exert deterministic effects on balance. This study examined whether low-amplitude nGVS (<1 mA), as typically used in stochastic resonance paradigms, directly influences postural behavior through stimulus-response coupling. Twenty healthy young adults stood on a force plate with feet together and eyes closed on either a rigid surface or 10-cm foam. In randomized order, they completed 300-second trials with band-limited (0-30 Hz), zero-mean nGVS at ±0, 0.1, 0.2, 0.3, 0.5, and 0.7 mA. Coupling between the stimulation waveform and mediolateral ground-reaction force was assessed using coherence and time-cumulant density. Mean coherence was significant mainly at higher amplitudes (0.5-0.7 mA) on both surfaces, whereas time-cumulant density identified significant time-locked vestibular-evoked response components at much lower amplitudes, down to 0.1 mA. These included an early response around 135-155 ms and a later, prominent response around 360-410 ms. Individually, significant coherence was common at 0.5-0.7 mA (15-19 of 20 participants), while cumulant-based responses appeared in some participants even at 0.1 mA. Responses were clearer on foam, consistent with greater vestibular reliance when somatosensory input is less reliable. Overall, low-amplitude nGVS can entrain postural output, suggesting that balance changes during “subsensory” stimulation may reflect both stochastic-resonance-like effects and deterministic vestibular drive, underscoring the need to quantify coupling alongside performance outcomes.

## INTRODUCTION

A contributor to falls in older adults and in patients with vestibular processing disorders is deterioration of the vestibular hair cells that sense motion, along with atrophy and desensitization of vestibular afferent nerve fibers that transmit motion information to the brain (Alvarez et al. 2000; Merchant et al. 2000; Richter 1980). These vestibular impairments can cause dizziness and loss of stability that could contribute to life-altering falls (Agrawal et al. 2009; Iwasaki and Yamasoba 2015; Politi et al. 2022). The consequences of falls are costly, both economically and socially, including increased hospitalization, health care, and nursing home costs, as well as fear of falling, functional decline, reduced mobility, depression, and greater burdens on families and society. (Clegg et al. 2013; Florence et al. 2018; Gale et al. 2016; Guinand et al. 2012).

The frequency and severity of fall-related outcomes motivate interventions that specifically target balance deficits linked to vestibular impairment. Recent evidence suggests that a sensory intervention known as noisy galvanic vestibular stimulation (nGVS) may reduce fall risk in older adults and in those with impaired vestibular information processing (Fujimoto et al. 2016; Inukai et al. 2018; Iwasaki et al. 2014; Wuehr et al. 2016, 2017). nGVS is the transcutaneous application of low-amplitude electrical noise to vestibular afferents via electrodes placed over the mastoid processes behind each ear and is a variant of the broader intervention GVS (Dlugaiczyk et al. 2019, 2020; Fitzpatrick and Day 2004; Lajoie et al. 2021; Marchand et al. 2025; McLaren et al. 2023; Stefani et al. 2020; Valter et al. 2025). Because many vestibular disorders involve reduced sensitivity to motion that contributes to postural instability, interventions that enhance vestibular motion sensitivity may lessen the impact of these impairments (Iwasaki and Yamasoba 2015; Mulavara et al. 2011).

nGVS is commonly framed as a “subsensory” stimulus: amplitudes are often selected to be imperceptible (either via sensation at the skin or induced motion perceptions) and behavioral changes are frequently interpreted through stochastic resonance (SR), wherein added noise can improve detection of otherwise weak signals (Fujimoto et al. 2016; Goel et al. 2015; Inukai et al. 2018; Iwasaki et al. 2014; McDonnell and Ward 2011; Moss 2004; Mulavara et al. 2015; Nooristani et al. 2019; Pal et al. 2010; Wuehr et al. 2016). Importantly, however, all forms of GVS directly target the nearby vestibular afferents (Goldberg et al. 1984; Kim and Curthoys 2004; Kwan et al. 2019) and therefor an “imperceptible” current may not be functionally subthreshold for vestibular pathways and may still exert a deterministic influence on motor output (Simoneau et al. 2025). This raises the possibility that the same low amplitudes used to probe SR may entrain postural behavior, producing responses qualitatively similar to those observed with suprathreshold stimulation (Dakin et al. 2007, 2010) but reduced in magnitude, thereby complicating the mechanistic interpretation of performance changes during nGVS.

Here our single aim was to determine whether the same low stimulation amplitudes (< 1mA) commonly used to probe SR also directly influence sway behavior via stimulus-response coupling. Specifically, we examined coherence and the time-cumulant density between the applied stimulus and medio-lateral horizontal forces while standing. Evidence of coupling would indicate that low-amplitude vestibular stimulation can entrain postural behavior, producing responses similar to those evoked by more generic GVS at higher amplitudes, but with smaller magnitude, and would therefore represent a confound when interpreting SR-like behavioral changes purely as improved inertial motion signal detectability by the vestibular pathways.

## METHODS

### Participants

Twenty healthy young adults (18–27 years; 22.2 ± 2.8 years; 171.4 ± 9.8 cm; 68.4 ± 14.9 kg; 12 females; mean ± SD) completed pre-screening health questionnaires (Physical activity readiness questionnaire and an electric vestibular stimulation screening questionnaire), COVID-19 protocol statements, and informed consent using HIPAA compliant REDCap software. Individuals were excluded if they reported recent lower-extremity injury, chronic sensory dysfunction (neurologic, vestibular, or visual), or an inability to stand for >15 min. All procedures were approved by the Utah State University Institutional Review Board (Protocol #10700).

### Experimental setup and stimulation

Participants stood barefoot with feet together completed two conditions, standing on a force-plate (off-foam) and standing on a block of foam on the force-plate (on-foam). Foot position was marked with tape to standardize placement across trials. In the foam condition, participants stood on a 10-cm-thick foam pad placed atop the force plate. For all trials, participants kept their arms at their sides and their eyes closed.

Before electrode placement, participants removed earrings and secured long hair. The skin over both mastoid processes was cleaned with an alcohol swab. Carbon rubber electrodes (~9 cm^2^) coated with Spectra 360 electrode gel were placed bilaterally over the mastoids using medical adhesive tape in a bipolar, binaural configuration.

The nGVS stimulus consisted of zero-mean, band-limited (0–30 Hz) white-noise current (Dakin et al. 2007; McLaren et al. 2023; Mulavara et al. 2011) generated in LabVIEW (National Instruments, Austin, TX, USA) and delivered via a digital-to-analog converter (PXIe-6363, National Instruments, Austin, TX, USA) to an isolated constant-current stimulator (STMISOL, Biopac, Goleta, CA, USA). Force-plate signals and the stimulation waveform were recorded using the same LabVIEW program at 2000 Hz and saved as text files.

### Experimental procedure

Participants completed 12 trials (six on the rigid force-plate surface and six on foam), with each trial lasting 300 s. After pseudo-randomly selecting the starting condition, trials alternated between the off-foam and on-foam conditions to limit fatigue associated with prolonged standing on foam. Within each surface condition, one of six nGVS amplitudes was applied: ±0, 0.1, 0.2, 0.3, 0.5, and 0.7 mA. The 0.4 mA and 0.6 mA conditions were omitted to reduce total trial count and participant fatigue. The six amplitudes were selected as this range is commonly used to elicit stochastic resonance-like behavior. Stimulation amplitudes were delivered in a randomized order, and participants were blinded to stimulation amplitude. Between trials, participants sat and rested for 2 min.

### Data analysis

Data were processed offline in MATLAB (R2021a; MathWorks, Natick, MA, USA) using custom scripts. To assess whether the vestibular stimulus drove postural responses, we calculated coherence and the time-cumulant density between the nGVS waveform and mediolateral ground-reaction force (Dakin et al. 2007, 2010). Coherence quantifies the strength of the linear relationship between two signals as a function of frequency; significant coherence indicates a consistent phase relationship between input stimulus frequencies and the same frequencies in the response (forces acting at the feet). The time-cumulant density functions as an unnormalized cross-correlation, showing how the two signals are related across different time lags.

Prior to coherence and time-cumulant density estimation, force-plate signals were low-pass filtered at 35 Hz using a zero-phase shift filter. Coherence and time-cumulant density were estimated for each participant using the NeuroSpec toolbox (Halliday et al. 1995)). Analyses used the full 300 s of data from each trial with a window length of 4096 samples (2.048 s at 2000 Hz), yielding 146 segments per trial and a frequency resolution of 0.4883 Hz (2000/4096). The 95% confidence limit for coherence was computed from the number of segments (Halliday et al. 1995).

For the mean time-cumulant density (**Figure 1, column 2**), each non-zero amplitude stimulation condition was compared against the 0 µA condition using non-parametric bootstrapped 95% confidence intervals (10000 bootstrap samples) at each time lag. Time regions were considered different when the confidence intervals did not overlap (i.e., the lower bound of one condition exceeded the upper bound of the other). In single subjects, significant peaks in the time-cumulant density were identified as those that exceeded the 95% confidence interval provided by the Neurospec library (Halliday et al. 1995). Both coherence and time-cumulant density were evaluated because components of the response may be significant in the cumulant density even when coherence does not reliably reach significance.

**Fig 1.**
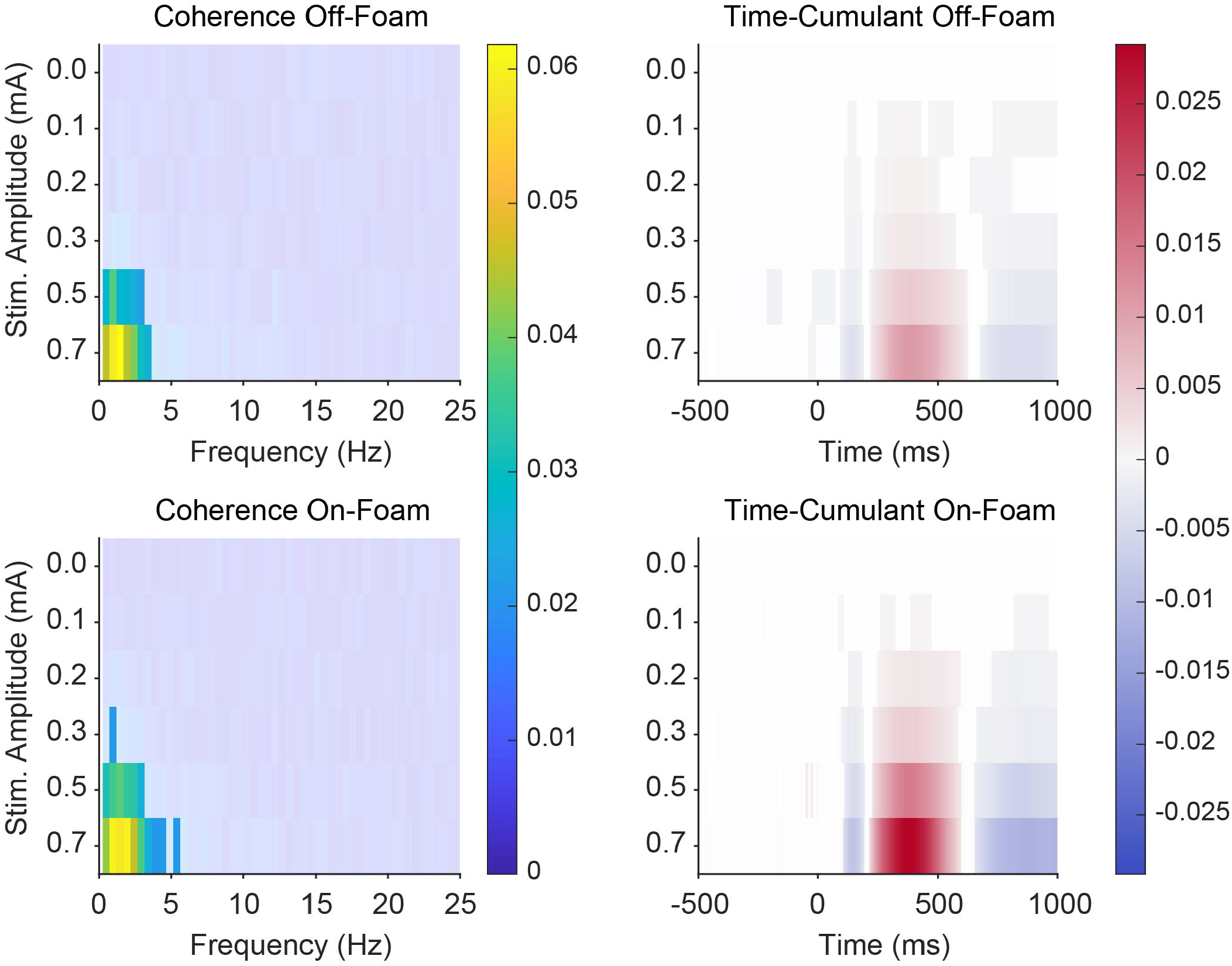
Significant Mean Coherence and Time-Cumulant Density. Mean coherence is shown in column 1. Vibrant colored regions indicated where mean coherence exceeded the confidence limit (0.0204), with the scale represented on the color bar to the right of the column. Significant regions for the time-cumulant density are shown in columns 2. Significance was established as regions where the bootstrapped confidence intervals for each stimulation trial exceeded the confidence interval for the 0mA condition. Blue indicates a negative waveform and red a positive waveform. The colormap displays near zero significant time-cumulant density values as light grey to contrast them with the white non-significant regions. Coherence is a unitless measure, and the time-cumulant density has units of NmA. Stim Amplitude is stimulus amplitude and is plotted in mA.

To document the prevalence of responses across individual participants and amplitudes, coherence spectra that exceed NeuroSpec’s coherence confidence limit for more than one consecutive point, or exceeded NeuroSpec’s time-cumulant density confidence interval, were recorded as presenting significant responses in each measure respectively. Once tabulated, responses that were not consistently present at sequentially increasing amplitudes were removed from the tabulation (e.g., 0.5 and 0.7 mA or 0.4, 0.5 and 0.7 mA were included, but significant responses observed at 0.1mA and not 0.2 mA were removed) (**Table 1**).

**Table 1.** Consistency of Single-Participant Responses. This table lists the number of participants in whom significant responses were consistently observed across stimulus amplitudes. Participants were counted only if significant responses were observed at the listed amplitude and at every subsequent higher stimulus amplitude. Stimulus amplitudes (mA) are shown on the top row of each chart. The time-cumulant density results are organized by response component.

|  |  | Coherence |  |  |  |  |  |  |  |  |  |  |  |
| --- | --- | --- | --- | --- | --- | --- | --- | --- | --- | --- | --- | --- | --- |
|  |  | Off-Foam |  |  |  |  |  | On-Foam |  |  |  |  |  |
| Stim. Amplitude (mA) |  | 0 | 0.1 | 0.2 | 0.3 | 0.5 | 0.7 | 0 | 0.1 | 0.2 | 0.3 | 0.5 | 0.7 |
| n |  | 0 | 0 | 1 | 5 | 15 | 18 | 0 | 0 | 2 | 6 | 17 | 19 |

**Table 1. Consistency of Single-Participant Responses.**
|  |  | Time-Cumulant Density |  |  |  |  |  |  |  |  |  |  |  |
| --- | --- | --- | --- | --- | --- | --- | --- | --- | --- | --- | --- | --- | --- |
|  |  | Off-Foam |  |  |  |  |  | On-Foam |  |  |  |  |  |
| Stim. Amplitude (mA) |  | 0 | 0.1 | 0.2 | 0.3 | 0.5 | 0.7 | 0 | 0.1 | 0.2 | 0.3 | 0.5 | 0.7 |
| Early Peak (n) |  | 0 | 0 | 1 | 2 | 7 | 13 | 0 | 0 | 1 | 3 | 7 | 14 |
| Second Peak (n) |  | 0 | 5 | 8 | 19 | 20 | 20 | 0 | 3 | 12 | 17 | 20 | 20 |
| Third Peak (n) |  | 0 | 1 | 2 | 3 | 11 | 14 | 0 | 2 | 8 | 9 | 13 | 18 |

## RESULTS

### General Results

All participants maintained standing balance throughout all 12 trials. However, participants reported that standing on the foam with feet together and eyes closed was more difficult. Anecdotally, five participants reported perceiving the stimulus during the 0.5 and 0.7mA trials, and one participant reported perceiving stimulation on every trial. Two participants reported worse balance during the 0.7 mA trial, while two participants reported perceiving improved balance during the 0.2 mA and 0 mA trials. Participants were not informed of the stimulus amplitudes, and no adverse effects were reported.

### Specific Results

The root mean square (RMS) stimulus amplitudes for the *off-foam* condition were: 0 mA = 1.1 µA RMS, 0.1 mA = 23.2 µA, 0.2 mA = 45.5 µA, 0.3 mA = 70.1 µA, 0.5 mA = 113.9 µA, and 0.7 mA = 163.3 µA. For the *on-foam* condition, RMS stimulus amplitudes were: 0 mA = 0.73 µA, 0.1 mA = 23.3 µA, 0.2 mA = 46.3 µA, 0.3 mA = 69.5 µA, 0.5 mA = 116.7 µA, and 0.7 mA = 161.0 µA.

Significant mean coherence was observed at 0.5 and 0.7 mA in both the off-foam and on-foam conditions (**Figure 1**), and in individual participants (**Table 1**). At the individual-participant level, significant coherence was observed in the off-foam condition for 0.2 mA (n = 1), 0.3 mA (n = 5), 0.5 mA (n = 15), and 0.7 mA (n = 18), and in the on-foam condition for 0.2 mA (n = 2), 0.3 mA (n = 6), 0.5 mA (n = 17), and 0.7 mA (n = 19).

In the time-cumulant density, significant responses were observed at amplitudes as low as 0.1 mA for the early negative response, the second positive peak, and the third negative peak. At the individual-participant level (**Table 1**), consistent significant early responses were observed in one participant starting at 0.2 mA in both off-foam and on-foam conditions. For the second and third peaks, consistent significant responses were observed starting at 0.1 mA, particularly for the second peak (off-foam: n = 5; on-foam: n = 3).

For the *off-foam* condition, the mean timing of the early response across participants was: 137.5 ms at 0.2 mA (n = 1), 137.5 ± 9.9 ms at 0.3 mA (n = 2), 141.4 ± 12.6 ms at 0.5 mA (n = 7), 140.9 ± 14.9 ms at 0.7 mA (n = 13). The mean timing of the second response in the *off-foam* condition was: 377.9 ± 53.7 ms at 0.1 mA (n = 5), 391.9 ± 31.9 ms at 0.2 mA (n = 8), 403.6 ± 63.9 ms at 0.3 mA (n = 19), 386 ± 47.6 ms at 0.5 mA (n = 20), 400.7 ± 37.4 ms at 0.7 mA (n = 20). We did not estimate the peak timing of the later response because it *often* extended beyond the analysis window.

For the *on-foam* condition, the mean timing of the early response across participants was: 134.5 ms at 0.2 mA (n = 1), 150 ± 23.9 ms at 0.3 mA (n = 3), 154.4 ± 15.1 ms at 0.5 mA (n = 7), 152.1 ± 16.6 ms at 0.7 mA (n = 14). The mean timing of the second response in the *on-foam* condition was: 358 ± 101.9 ms at 0.1 mA (n = 3), 409.3 ± 73 ms at 0.2 mA (n = 12), 375.5 ± 44.7 ms at 0.3 mA (n = 17), 393.6 ± 45.1 ms at 0.5 mA (n = 20), 370.2 ± 39.8 ms at 0.7 mA (n = 20).

*Selected i*ndividual participant responses are shown in ***F*igure 2**. Columns 1-2 illustrate two examples of *higher performing* participants with more prominent coherence, while columns 3-4 illustrate two examples of *lower performing* participants with weaker coherence. The time-cumulant density plots follow the same layout: columns 1 and 3 show participants with more prominent responses at lower amplitudes, whereas columns 2 and 4 show participants with weaker responses. Generally, there was significant between subject variance in the prominence with which coupling could be observed between participants.

**Fig 2.**
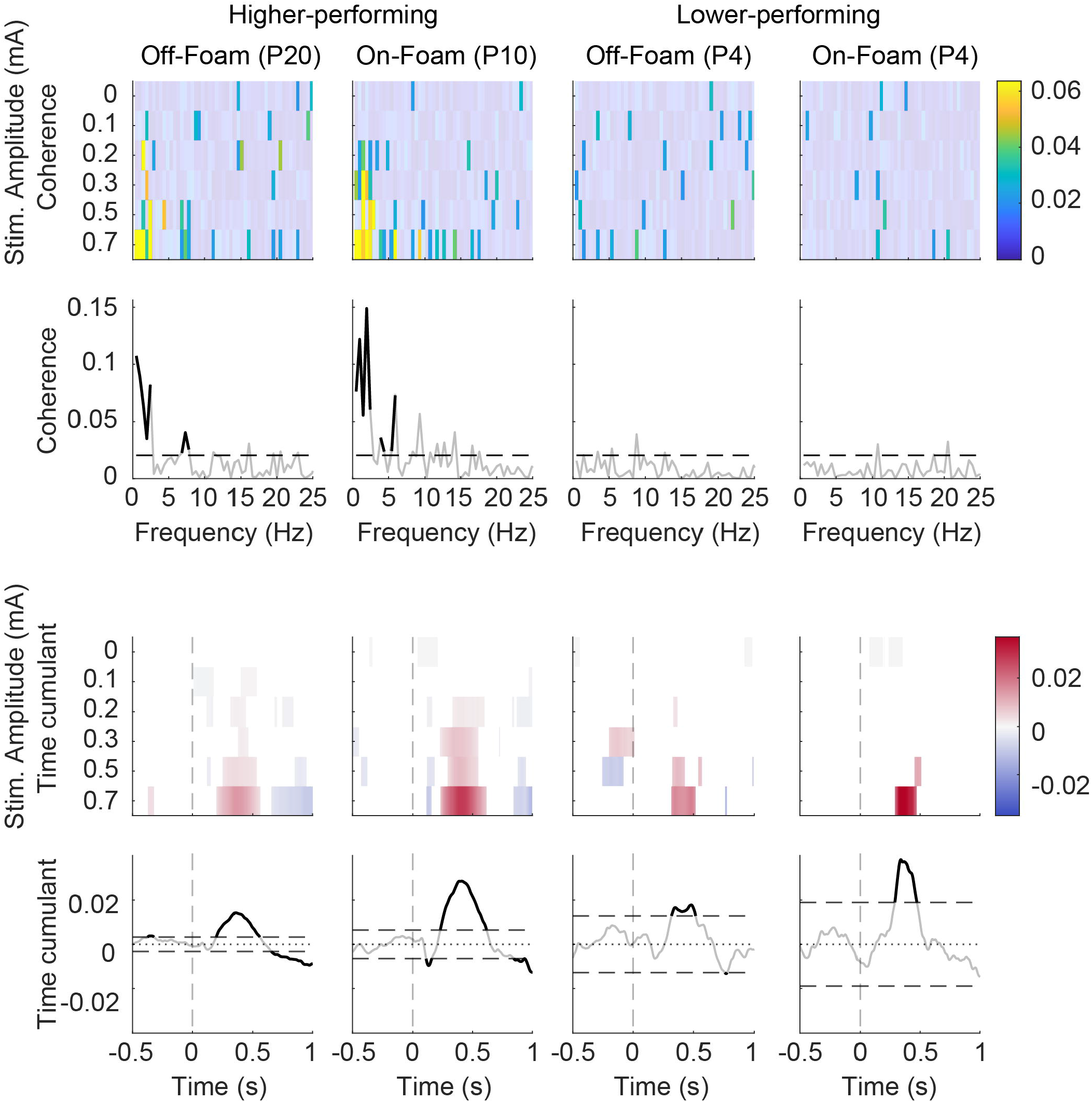
Example individual-participant coherence and time-cumulant densities for higher- and lower-performing participants. Row 1 shows coherence across stimulus amplitudes, with coherence magnitude indicated by color; values below Neurospec’s confidence limit are transparent. Row 2 shows the coherence spectra for the 0.7 mA condition, non-significant regions are transparent, and the Neurospec library’s confidence limit indicated with a dashed line. Row 3 shows heat maps of significant peaks in the time-cumulant density, based on Neurospec’s confidence interval; non-significant regions are white. Row 4 shows the time-cumulant density for the 0.7 mA condition, with confidence intervals indicated by the large dashed horizontal lines; non-significant regions are transparent. Zero is indicated by the faint dotted line. The left columns show individual-subject data from participants with stronger stimulus-evoked responses, whereas the right columns show participants with weaker responses. In the higher-performing participants, significant responses with appropriate timing are sometimes evident at stimulus amplitudes as low as 0.1 and 0.2 mA. In contrast, although some significant regions are present in the lower-performing participants, their time-cumulant density estimates exhibit much wider confidence intervals and significant peaks are observed only at the higher stimulus amplitudes. irence Off-Foam Time-Cumulant Off-Foam

## DISCUSSION

This study examined whether the low nGVS amplitudes commonly used in stochastic resonance (SR) paradigms (Eder et al. 2022; Iwasaki et al. 2014; Mulavara et al. 2011, 2015; Stefani et al. 2020) can directly drive postural behavior through stimulus–response coupling. Importantly, detecting a stimulus-locked coupling component does not, by itself, establish that nGVS produces a practically meaningful change in whole-body stability; the functional relevance of coupling ultimately depends on how its magnitude relates to sway or stability outcomes. Across both rigid and foam surfaces, coupling was detected, but the capacity to resolve it depended on the analysis method: mean coherence reached significance mainly at higher amplitudes (0.5-0.7 mA), while the time-cumulant density revealed significant, time-locked response components at much lower amplitudes (down to 0.1 mA). This divergence suggests that low-amplitude nGVS produces vestibular-evoked responses at the feet but that the coupling is not sufficiently strong or spectrally concentrated to produce significant coherence below 0.5 mA. Responses were generally more prominent on foam, which was anticipated, potentially due to reduced reliability or predictability of somatosensory cues and associated sensory reweighting (Cenciarini and Peterka 2006; Ernst 2006; Peterka 2002). This raises the possibility that in contexts that increase reliance on vestibular cues (e.g., compliant surfaces and/or reduced visual input), stimulus-locked drive, and therefore its potential confounding influence on SR-like interpretations, could be amplified along with possible beneficial effects of SR. In older adults and clinical populations, where multisensory reliability and weighting may differ from young healthy adults, the magnitude and behavioral consequences of such coupling remains unclear.

These findings may have important implications for mechanistic interpretation of “SR-like” effects (McDonnell and Ward 2011; Stefani et al. 2020). Because galvanic stimulation directly modulates vestibular afferent activity (Curthoys and MacDougall 2012; Dlugaiczyk et al. 2020; Goldberg et al. 1984; Kim and Curthoys 2004; Kwan et al. 2019), the observation of stimulus-locked force responses at low amplitudes indicates that “imperceptible” current may not be functionally subthreshold for vestibular-motor pathways (Simoneau et al. 2025). In other words, performance changes during low-amplitude nGVS could reflect a mixture of (i) true SR-related improvements in processing weak endogenous vestibular signals (McDonnell and Ward 2011; Moss 2004; Yamamoto et al. 2005) and (ii) deterministic entrainment of postural output by the applied stimulus (Britton et al. 1993; Coats and Stoltz 1969; Lund and Broberg 1983; Pavlik et al. 1999; Scinicariello et al. 2003; Simoneau et al. 2025). Therefore, changes in balance metrics during nGVS should not be attributed to SR alone without quantifying and ideally accounting for the effect of stimulus–response coupling. In practice, coupling estimates could be incorporated as covariates or used to partition stimulus-locked versus non-locked variance when interpreting balance outcomes.

A related possibility is that even a mild change in vestibular drive could alter the dynamics of stance. From a feedback-control perspective, nGVS may inject an exogenous vestibular-like signal into the sensory estimate used to generate corrective torques. Thus, even when perceptually “subsensory,” any coherent vestibular component could bias state estimation and subtly shift effective feedback gains or sway frequency content (Cenciarini and Peterka 2006; Fetsch et al. 2009; Peterka 2002; Peterka and Loughlin 2004). Consistent with this view, vestibular-evoked balance responses to stochastic/multisine stimulation can be described with an approximately linear input–output mapping (Forbes et al. 2014; Hannan et al. 2021) over relevant operating ranges, while response magnitude and time course depend on task context and the reliability of other sensory cues (Blouin et al. 2011; Day and Guerraz 2007; Ernst 2006; Héroux et al. 2015; Horslen et al. 2014). Accordingly, low-amplitude nGVS may act not only as “noise” but also as a mild vestibular perturbation that can alter postural dynamics, reinforcing the need to quantify coupling when interpreting changes in balance outcomes.

The latency structure of the time-cumulant density responses supports the interpretation that low-amplitude stimulation is inducing established vestibular-evoked postural dynamics that are well described in suprathreshold GVS/SVS work (Britton et al. 1993; Cathers et al. 2005; Dakin et al. 2007, 2010; Fitzpatrick et al. 1994; Fitzpatrick and Day 2004; Mian et al. 2010). In our data, the early component occurred around ~135–155 ms, and the later prominent component occurred around ~360–410 ms. Prior work using vestibular stimulation and shear ground-reaction forces reports short- and medium-latency force peaks on the order of ~120–200 ms and ~290–400 ms, respectively, with the force timing reflecting electromechanical delay relative to earlier EMG onsets (Britton et al. 1993; Dakin et al. 2010; De Melker Worms et al. 2017; Fitzpatrick et al. 1994; Horslen et al. 2014; Mian et al. 2010). While our second peak tended to be somewhat later than some stochastic vestibular stimulation - ground reaction force reports, later portions of the response (approaching and exceeding ~400 ms) have been argued to incorporate greater multisensory feedback contributions depending on task context, which could contribute to timing differences across protocols and postural constraints (e.g., foam, eyes closed, feet together).

Finally, we observed substantial inter-individual variability, with clear ‘strong’ and ‘weak’ responders, suggesting that a given amplitude should not be assumed to be uniformly ‘subsensory’ across participants, even though sensory thresholds are typically calibrated individually. Moreover, because this study used healthy young adults, the results primarily show that low-amplitude nGVS can produce measurable coupling in an intact system and that, in this population, its effects are not clearly distinguishable from those of higher amplitude noisy or stochastic galvanic vestibular stimulation (> 1mA). However, generalization to older adults or vestibular-impaired groups remains to be tested. Undoubtedly, a limitation of this study is not collecting different measures of thresholds (and impedances) to determine how well the threshold amplitudes correlate with the observation of nGVS-force response coupling in individual subjects. Future work could combine individualized thresholding with full-body kinematics or center-of-pressure outcomes, and directly evaluate whether the magnitude of stimulus - response coupling predicts changes in stability (benefit vs disruption), especially in the clinical populations most relevant to fall risk.

## Conclusion

The low-amplitude nGVS in the range 0.1 mA - 0.7 mA produces measurable stimulus–response coupling between the stimulation waveform and mediolateral ground-reaction forces, with time-cumulant density detecting time-locked vestibular-evoked components even at 0.1 mA (and coherence becoming prominent mainly at 0.5–0.7 mA). Coupling tended to be more evident on foam and varied substantially across individuals, implying that balance or behavioral changes during low-amplitude nGVS may reflect direct vestibular modulation and deterministic entrainment in addition to (or instead of) stochastic resonance-like enhancement of endogenous vestibular signals.

